# Familiarity of Background Music Modulates the Cortical Tracking of Target Speech at the Cocktail Party

**DOI:** 10.1101/2022.07.14.500126

**Authors:** Jane A. Brown, Gavin M. Bidelman

**Author notes:** **Address for editorial correspondence:** Gavin M. Bidelman, PhD, Department of Speech, Language and Hearing Sciences, 2631 East Discovery Parkway, Bloomington, IN 47408.

## Abstract

The “cocktail party” problem – how a listener perceives speech in noisy environments – is typically studied using speech (multi-talker babble) or noise maskers. However, realistic cocktail party scenarios often include background music (e.g., coffee shops, concerts). Studies investigating music’s effects on concurrent speech perception have predominantly used highly controlled synthetic music or shaped noise which do not reflect naturalistic listening environments. Behaviorally, familiar background music and songs with vocals/lyrics inhibit concurrent speech recognition. Here, we investigated the neural bases of these effects. While recording multichannel EEG, participants listened to an audiobook while popular songs (or silence) played in the background at 0 dB signal-to-noise ratio. Songs were either familiar or unfamiliar to listeners and featured either vocals or isolated instrumentals from the original audio recordings. Comprehension questions probed task engagement. We used temporal response functions (TRFs) to isolate cortical tracking to the target speech envelope and analyzed neural responses around 100 ms (i.e., auditory N1 wave). We found that speech comprehension was, expectedly, impaired during background music(s) compared to silence. Target speech tracking was further hindered by the presence of vocals. When masked by familiar music, response latencies to speech were less susceptible to informational masking, suggesting concurrent neural tracking of speech was easier during music known to the listener. These differential effects of music familiarity were further exacerbated in listeners with less musical ability. Our neuroimaging results and their dependence on listening skills are consistent with early attentional gain mechanisms where familiar music is easier to tune out (listeners already know the song’s expectancies) and thus can allocate fewer attentional resources to the background music to better monitor concurrent speech material.

## INTRODUCTION

Listeners are constantly faced with the challenge of listening to speech in noisy environments. This so-called “cocktail party” problem is often studied using noise or multi-talker babble maskers. However, many realistic cocktail party scenarios also involve music (e.g., coffee shops, concerts), which is not often considered in studies of auditory scene analysis. The effect of background music on concurrent speech/linguistic tasks is mixed and dependent on many factors, including type of music, participant characteristics, and task structure (reviewed by (Kämpfe et al., 2011). Music presented concurrently with a memorization task impairs performance, but only for listeners who prefer to study without background music (Crawford & Strapp, 1994). Fast-tempo background music increases spatial processing speed and linguistic processing accuracy (Angel et al., 2010), but at the same time, can disrupt reading comprehension (Thompson et al., 2011). However, it is clear that music with vocals more negatively affects concurrent tasks across various cognitive modalities (Brown & Bidelman, 2022; Crawford & Strapp, 1994; Darrow et al., 2006; Lee et al., 2020; Martin et al., 1988; Perham & Currie, 2014; Vasilev et al., 2018). Linguistic content in the masker introduces informational masking, which in turn interferes with cognitive resources needed to complete the task.

In typical (i.e., speech-on-speech) cocktail party tasks, the familiarity of a talker can be advantageous for speech recognition (Souza et al., 2013; Yonan & Sommers, 2000) when the familiar voice is either the target or the masker (Johnsrude et al., 2013). A familiar voice is retained implicitly (Yonan & Sommers, 2000), which allows for more efficient processing of novel words or sentences spoken in that voice (Pisoni, 1993). However, the role of familiarity for music in perceptual-cognitive tasks is not well known. It is also worth noting that many studies define “familiarity” differently (e.g., exposure training of naïve listeners versus songs that listeners already know), which limits comparisons between results. Still, familiar music maskers can improve various linguistic behavioral measures (Feng & Bidelman, 2015; Russo & Pichora-Fuller, 2008), but may be detrimental to foreign language learning (De Groot & Smedinga, 2014). Previous work from our lab (Brown & Bidelman, 2022) has shown a negative familiarity effect of background noise on concurrent speech recognition. In a music-on-speech cocktail party task, we found speech recognition performance was worse during familiar compared to unfamiliar music maskers, likely due to the increased cognitive load of the familiar music (i.e., those songs were more distracting). However, that prior work was solely behavioral and did not provide insight into the neural underpinnings of those perceptual-cognitive effects.

Besides indexical attributes of the signal, demographic properties of the *listener* also modulate cocktail party perception (Bidelman & Dexter, 2015). In particular, musicality has been widely shown to alter auditory-cognitive brain structure and function, providing a “musician advantage” in various listening skills (Strait & Kraus, 2011). This is especially evident in speech-in-noise tasks where musicians show better degraded speech perception and more successful suppression of acoustic distractors (Bidelman & Krishnan, 2010; Bidelman & Yoo, 2020; Coffey et al., 2017; Parbery-Clark, Skoe, & Kraus, 2009; Yoo & Bidelman, 2019). Musicians’ improved speech-in-noise abilities might result from their superiority juggling multiple auditory streams (Bidelman & Yoo, 2020; Zendel & Alain, 2009) and lesser susceptibility to informational masking than their non-musician peers (Oxenham et al., 2003; Yoo & Bidelman, 2019). Musicians have more experience with auditory stream segregation (e.g., parsing a melody from harmonies, hearing one’s own melody in an orchestra), which in turn seems to enhance the parsing of degraded speech (Parbery-Clark, Skoe, Lam, et al., 2009). However, in contrast to speech-on-speech, musicians are more affected by background music than non-musicians (Patston & Tippett, 2011). Importantly, the “musician advantage” for speech-in-noise listening is not dependent on formal musical training. Mankel and Bidelman (2018) demonstrated similar effects in highly musical people without formal musical training, indicating that superior cocktail party skills may be attributed more to general listening abilities rather than music experiences/training, *per se.*

Extending this prior work, the current study sought to further investigate the role of background music familiarity and presence of vocals on concurrent speech perception. In a variant of the cocktail party task, participants listened to an audiobook (speech) in the presence of popular music maskers that varied in their familiarity to the listener and content of the original audio recording (e.g., isolated vocal or instrumentals). Our design departs from previous studies investigating the effects of music on speech intelligibility which have predominantly used synthesized music or music-like stimuli (Ekström & Borg, 2011; Eskridge et al., 2012) which, though easier to control, are not as ecologically valid as the popular music used here. Participants answered comprehension questions about the story to ensure task engagement with target speech material. We simultaneously recorded multichannel EEG and measured neural tracking to the speech envelopes using temporal response functions (TRFs) (Crosse et al., 2016). In accordance with our previous behavioral work (Brown & Bidelman, 2022), we hypothesized that the perception and neural tracking of speech would diminish (i.e., lower comprehension scores, TRFs with weaker amplitudes and longer latencies) when concurrent background music was familiar to listeners and when it contained vocals. These findings would suggest that speech perception suffers from a concurrent linguistic masker even from a different domain (i.e., music), as well as stronger attentional (mis)allocation to background music when it is familiar to the listener.

## MATERIALS & METHODS

### Participants

The sample included N=17 young adults ages 21-32 (*M*=25, *SD*=2.6 years, 4 male). This sample size is comparable with previous studies investigating continuous speech processing with TRFs (Ding et al., 2014; Forte et al., 2017; Lalor et al., 2009). 12 participants reported having musical training (*M*=10.25 years, *SD*=5.1). All participants showed audiometric thresholds better than 20 dB HL (octave frequencies, 250-8000 Hz) and reported English as their native language. Listeners were primarily right-handed (mean 75% laterality using the Edinburgh Handedness Inventory (Oldfield, 1971)). Each was paid for their time and gave written informed consent in compliance with a protocol approved by the IRB at the University of Memphis.

### Stimuli & task

During EEG recording, we measured neural speech tracking and comprehension by presenting a continuous audiobook in different background music conditions. The audiobook (taken from librivox.org) was *Doctor* by Murray Leinster, read by a male speaker. Silences > 300 ms were shortened to decrease extended silence in the stimulus while still sounding like natural speech (Ding & Simon, 2012b). The speech signal was RMS amplitude normalized and separated into 20 successive 2-minute segments.

Music stimuli were a subset of those used in Brown and Bidelman (2022), which had been previously identified as being familiar (“Just Dance” by Lady Gaga”) or unfamiliar (“Play with Fire” by Hilary Duff) to a cohort of young, normal-hearing listeners. We used a machine learning algorithm trained to separate instrumental and vocal tracks (lalal.ai) to isolate the instrumentals from the full unprocessed song (Brown & Bidelman, 2022). All music files (two songs each with the full song and isolated instrumentals) were sampled at 44100 Hz, converted to mono for diotic presentation, and RMS amplitude normalized after silences to equate the sound level across clips.

For each participant, the 20 audiobook segments were presented sequentially, separated into 4 runs (totaling 8 minutes of listening per condition). Each run contained five audiobook segments presented with each song condition (familiar with vocals, familiar without vocals, unfamiliar with vocals, and unfamiliar without vocals) and in silence. Participants were instructed to listen to the audiobook and ignore the background music. Each trial was 2 min, after which participants answered one comprehension question presented on a computer screen (20 questions total across the full experiment).

After completing the EEG task, participants completed the Profile of Music Perception Skills (PROMS) (Law & Zentner, 2012) to measure their musical listening skills. We have previously shown that high-PROMS-scoring individuals (“musical sleepers”) have enhanced speech processing akin to trained musicians despite having no formal musical training (Mankel & Bidelman, 2018). The PROMS contains 8 subtests focusing on different musical domains (e.g., rhythm, melody, timbre, etc.). For each subtest, participants heard two tokens (e.g., two melodies) and indicated whether they were the same or different. The scores were on a 5-point Likert scale, where correctly identifying “definitely same/different” was given one point and “probably same/different” was worth one-half point. The maximum possible test score was 80 (8 subtests, 10 items each).

After completing the experiment proper, participants rated each song on a scale from 1 (“Not at all familiar”) to 5 (“Extremely familiar”) to gauge their prior familiarity with the music stimuli.

### EEG recording procedures

Participants were seated in an electrically shielded, sound-attenuated booth for the duration of the experiment. Continuous EEG recordings were obtained from 64 channels aligned in the 10-10 system (Oostenveld & Praamstra, 2001) and digitized using a sample rate of 500 Hz (SynAmps RT amplifiers; Compumedics Neuroscan, Charlotte, NC). Contact impedances were maintained <10 kΩ. Music and speech stimuli were each presented diotically at 70 dB SPL through electromagnetically-shielded ER-2 insert headphones (Etymotic Research), resulting in a signal-to-noise ratio of 0 dB. Stimulus presentation was controlled with a custom MATLAB program (MathWorks, Natick, MA) and routed through a TDT RZ6 digital signal processor (Tucker-Davis Technologies, Alachua, FL). EEGs were re-referenced to average mastoids for further analysis. Data from 0-1000 ms after the onset of each 2-minute epoch were discarded to avoid transient brain responses in the subsequent analysis (Crosse et al., 2021). Epochs were then concatenated for each of the five conditions, resulting in 8 min of continuous data per condition.

### Behavioral data analysis

We logged the comprehension question response at the end of each presentation. Questions were scored as a binary “correct” or “incorrect” label.

### Electrophysiological data analysis: Temporal response functions (TRFs)

We analyzed continuous neural tracking to the speech signal using the Temporal Response Function toolbox in MATLAB (Crosse et al., 2016). The TRF is a linear function representing the (deconvolved) impulse response to a continuous stimulus. To measure EEG tracking to the speech, we extracted the temporal envelope of the audiobook via the Hilbert transform. EEG data were down-sampled to 250 Hz, then filtered between 1 and 30 Hz to isolate cortical activity to the low-frequency speech envelope. EEG and stimulus signals were both z-score normalized. As with conventional event-related potentials (ERPs), TRFs were computed for each participant to account for inherent inter-subject variability in neural response tracking (Crosse et al., 2021). We used 6-fold cross-validation to derive TRFs per condition. Ridge regression (Kulasingham & Simon, 2022) was used to identify the optimal λ smoothing parameter of the forward model for the speech-only condition. Model tuning was conducted using the speech-only condition to optimize the TRF to the clean (unmasked) speech. We used a fronto-central channel cluster (F1, Fz, F2, FC1, FCz, FC2, C1, Cz, C2) to further optimize the model fit to canonical topography of auditory ERPs. For each participant, the optimal λ was taken as the ridge parameter yielding the highest reconstruction of simulated neural response (i.e., correlation r-value between actual EEG and TRF-derived responses). We then used each participant’s optimal λ parameter to derive TRFs for all other conditions. This approach preserves response consistency within subjects while avoiding model overfitting (Crosse et al., 2021). The resulting TRF waveforms represents the time-varying weights how the EEG signal at each electrode changes in response to a unit change in the speech stimulus envelope.

We analyzed TRFs (i.e., RMS amplitude and latency) between 100-150 ms corresponding to the “N1” wave of the canonical auditory ERPs (see **Figure 1**). The N1 was selected as it reflects the early arrival of sound information in auditory cortex and is also modulated by attention (Hillyard et al., 1973; Näätänen & Picton, 1987; Picton et al., 1978). Previous EEG studies have also demonstrated noise has the largest effect on speech TRFs within this time window (Muncke et al., 2022). To further investigate possible hemisphere differences, we created two homologous channel clusters over front right (Fz, F2, F4, F6, F8, FC6, FT8) and front left (Fz, F1, F3, F5, F7, FC5, FT7) scalp regions.

**Figure 1.**
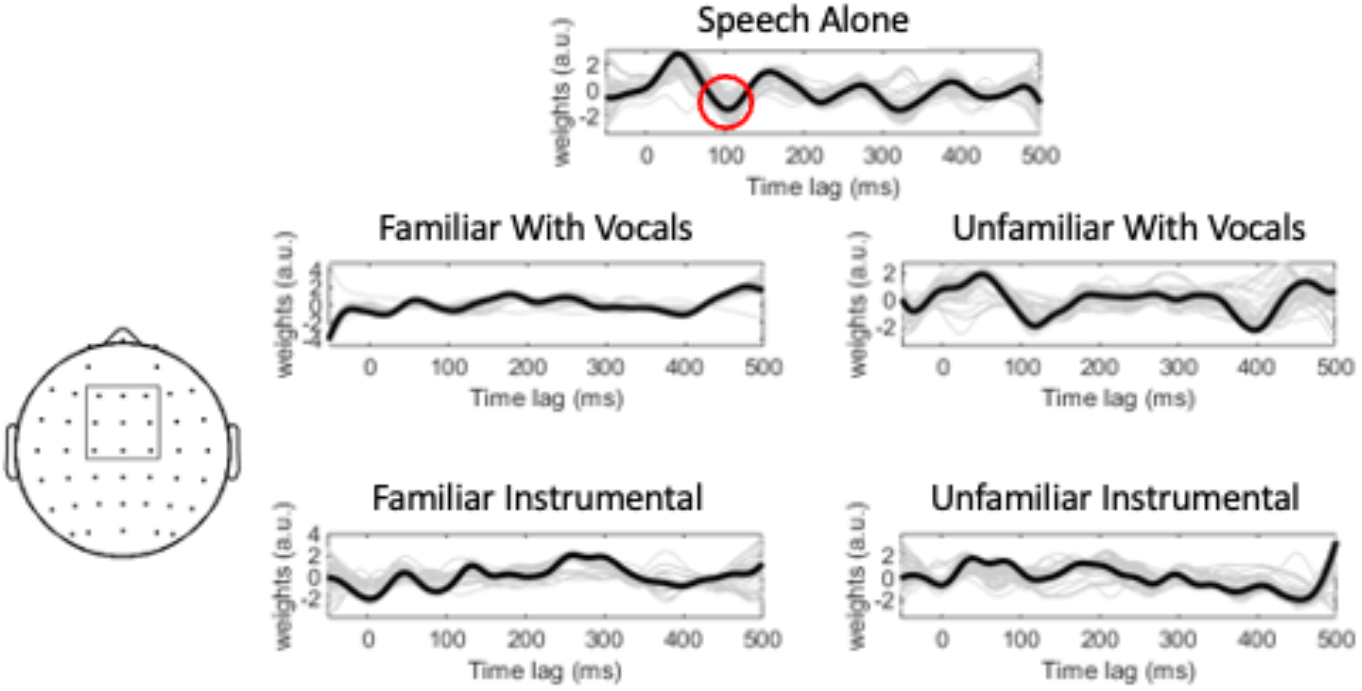
TRF waveforms (plotted at channel FCz) from a representative subject across music masker conditions. The model was trained using a fronto-central cluster of electrodes (left). Red circle = region of interest for analysis corresponding to the auditory N1. Bold black lines represent the TRF in each condition, and the grey lines show individual fits while training the model.

### Statistical analyses

Statistics were run in R using the lme4 package (Bates et al., 2015). For all analyses, we used mixed-effects models with fixed factors of familiarity (2 levels: familiar, unfamiliar) and song condition (2 levels; with vocals, without vocals). Subjects and trial served as random factors (where applicable). Because the behavioral response was a binary score (correct vs. incorrect), we analyzed those data using a generalized linear mixed-effects model ANOVA with binomial link function. Note that the Wald statistic is used to determine significant factor(s) effects for these models instead of conventional *F*-statistics (used in lme4 package; Bates et al., 2015). Peak amplitudes and latencies were normally distributed and thus analyzed using conventional linear mixed models and *F-*statistics. Multiple comparisons were corrected with Tukey adjustments. For all measures (score, latency, and amplitude), there were no hemisphere differences (all *ps* > 0.17), so subsequent analyses used data pooled between the two electrode clusters.

## RESULTS

### Behavioral data

Participants showed stark differences in their familiarity ratings across music selections (*t*(16)=19.13, *p*<0.001), validating our first stimulus manipulation. Speech comprehension scores were subject to a strong masking effect; comprehension was (expectedly) better for speech presented in silence compared to all other music-masked conditions (*t*(338)=3.23, *p*=0.001). An ANOVA based on the generalized binomial model showed no interaction between familiarity and song condition (χ^2^(1, N=17) = 1.26, *p*=0.26). There was no main effect of familiarity (χ^2^(1, N=17)=0.41, *p*=0.52). There was, however, a significant effect of song condition (χ^2^(1, N=17) = 5.42, *p*=0.02), whereby behavioral performance was overall poorer during vocal vs. non-vocal music maskers (**Figure 2**). These results confirm the effectiveness of music in masking target speech recognition as well as the added hinderance of music containing vocal (linguistic) information (cf. Brown & Bidelman, 2022).

**Figure 2.**
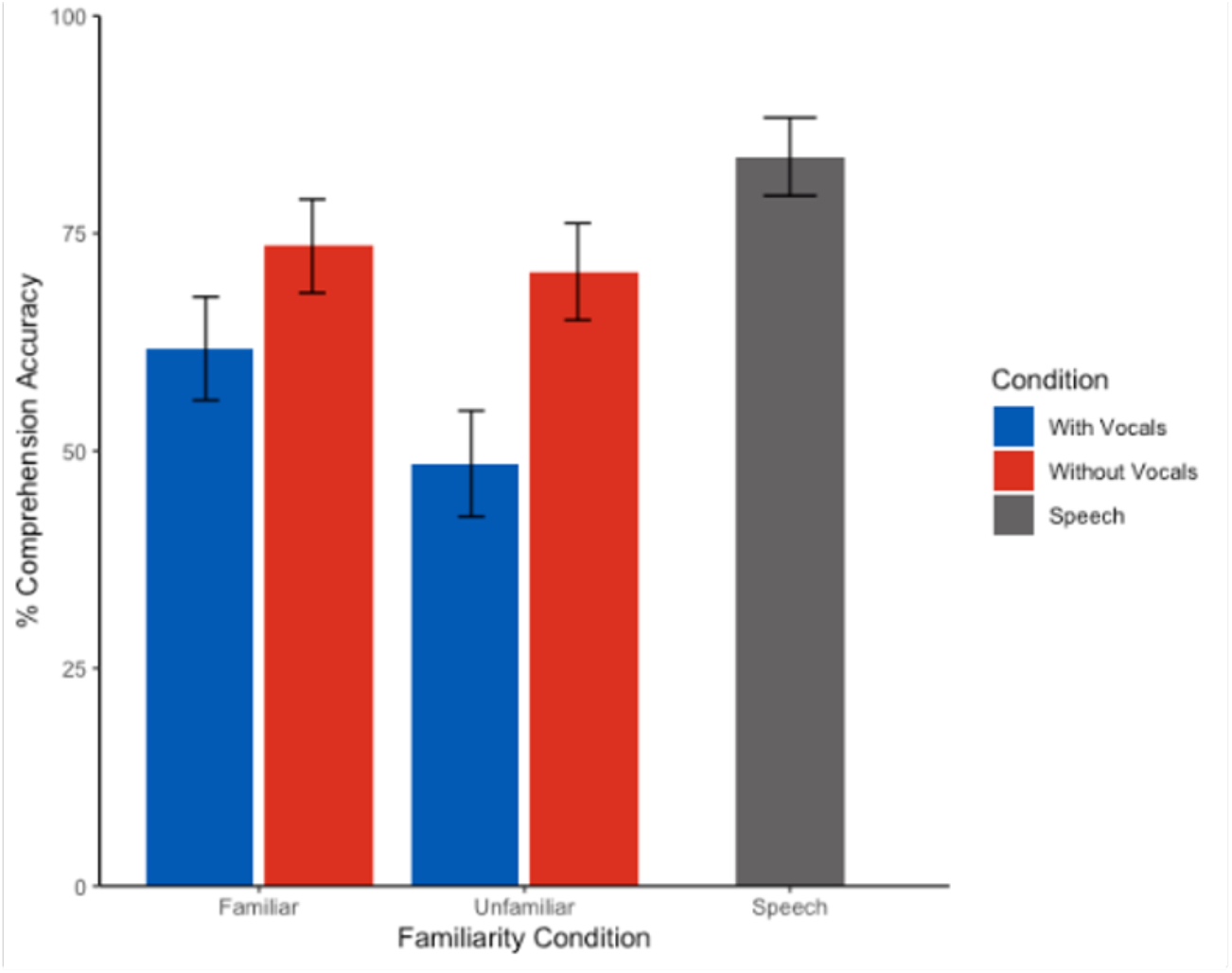
Speech comprehension scores during background music as a function of music familiarity and vocal condition. Speech recognition was poorer during concurrent vocal vs. non-vocal music. Error bars= ± 1 s.e.m.

### Electrophysiological data

TRF latency and amplitude are shown across conditions in **Figure 3**. Music-masked speech showed longer latencies than speech presented in silence (*t*(83)=3.40, *p*=0.001). An ANOVA conducted on TRF latencies revealed an interaction between familiarity and condition (*F*(1,48)=6.18, *p*=0.016, η^2^=0.11). Post hoc tests showed this interaction was attributable to longer speech-evoked TRF latencies in music with vocals than instrumentals alone (*t*(48)=2.54, *p*=0.015), but only in unfamiliar music. This vocal vs. instrumental latency differences was not observed for familiar music (*p*=0.32).

**Figure 3.**
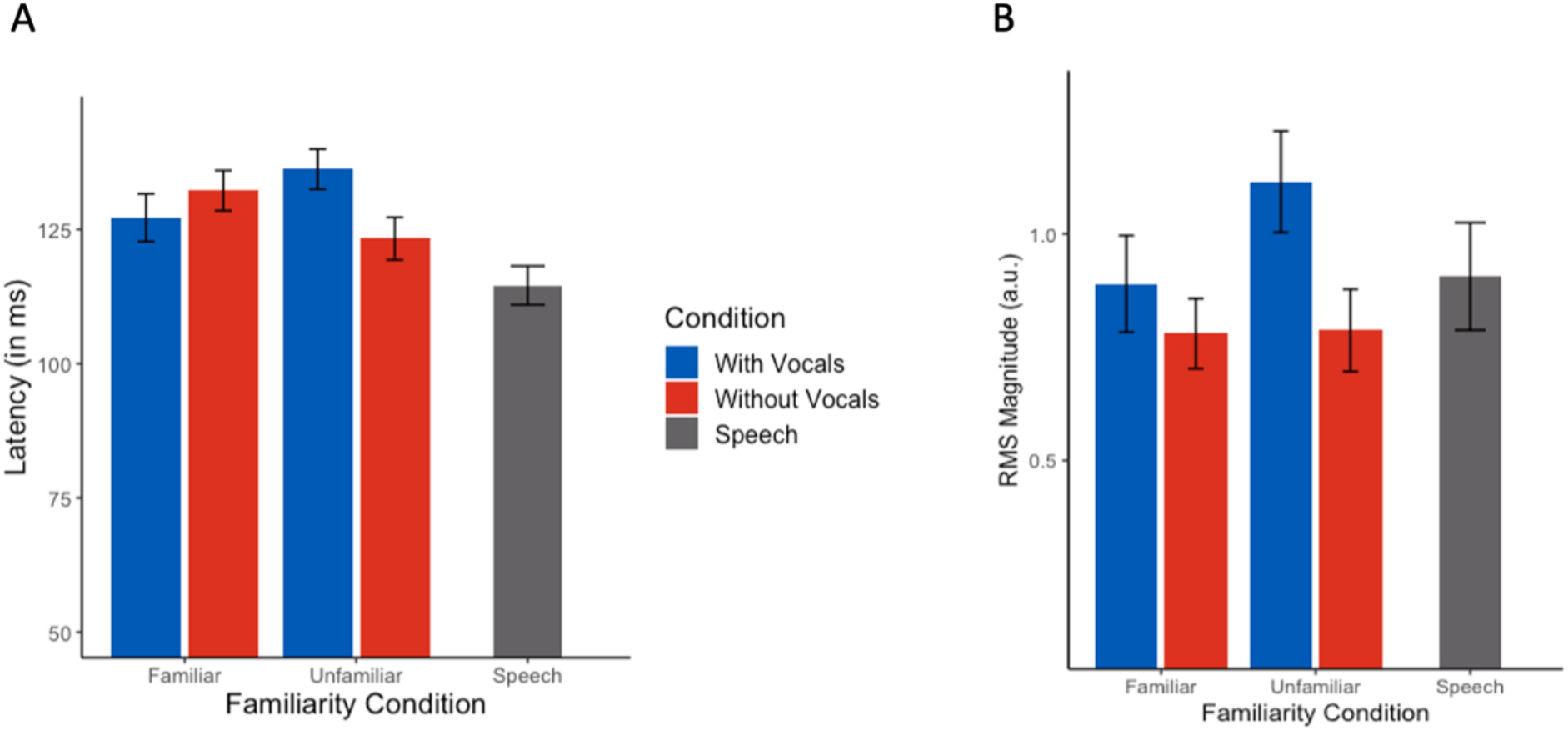
TRF N1 latencies and magnitudes across music masking conditions. *(A)* Latencies were longer for songs with vocals than instrumentals only during unfamiliar music. *(B)* Neural tracking of target speech was also stronger during music with vocals across the board. Error bars = ±1 s.e.m.

In contrast to latency measures, TRF amplitudes were less affected by masking and concurrent music (masking effect: *t*(83)=0.11, *p*=0.91). An ANOVA conducted on TRF RMS amplitudes, indicating the strength of speech tracking, showed a sole main effect of condition (*F*(1,48)=5.13, *p*=0.028, η^2^=0.10); responses were larger in music with vocals as compared to instrumentals. There was no effect of familiarity (*F*(1,48)=1.45, *p*=0.23) nor a condition*familiarity interaction (*F*(1,48)=1.28, *p*=0.26).

We ran Spearman’s correlations to determine the relationship between our electrophysiological and behavioral results. We found a negative association between TRF latency and speech comprehension scores (*r_s_*=-0.31, n=85, p=0.004) when aggregating data across all listeners and conditions. This indicates that longer TRF latencies corresponded with poorer speech comprehension. There was no correlation between behavior and TRF N1 amplitudes (*r_s_*=0.13, n=85, p=0.228). These data reveal the behavioral relevance of the TRF data: more delayed neural tracking of target speech is predictive of worse speech recognition performance.

### Neural speech tracking as a function of listeners’ musicality

We next asked whether speech-envelope tracking amidst music (as indexed via TRFs) varied as a function of listeners’ music-listening skills (as indexed by their PROMS scores). Previous studies demonstrate that individuals who lack formal music training but who nonetheless have superior auditory skills show advantages with speech identification and cocktail party processing (Mankel et al., 2020; Mankel & Bidelman, 2018). As in Mankel and Bidelman (2018), we divided our participants into two groups – “high PROMS” and “low PROMS” - using a median split of their PROMs musicality scores. The groups did not differ in age (t(15)=1.28, p=0.22), years of education (t(15)=0.82, p=0.43), or sex (Fisher’s exact test;*p*=0.294). The high PROMS group had 11.4 (*SD*=5.1) years of musical training as compared to the low PROMS group (*M*=3.6, *SD*=5.2 years; t(15)=3.13, p=0.01). To account for this difference in training, we ran our omnibus ANOVA models with the three factors of interest (familiarity, vocals condition, and PROMS level), and years of training as a covariate. In addition to dichotomizing the musicality variable (cf. MacCallum et al., 2002), we also ran models treating PROMS score as a continuous variable.

#### Behavior

We first tested for differences between PROMS scores and familiarity ratings to assess whether high vs. low musicality listeners were more/less familiar with the stimuli used in our experiment. There was no group difference for either the familiar (*t*(15)=1.63, *p*=0.125) or unfamiliar (*t*(15)=1.24, *p*=0.236) stimuli, indicating listeners were similarly (un)familiar with our song selections.

Analysis of speech comprehension during the EEG task revealed a significant condition*group interaction (χ^2^(1) = 4.12, *p*=0.042), as well as a marginal familiarity*condition*group interaction (χ^2^(1) = 3.63, *p*=0.056) (**Figure 4**). Group differences were also partially driven by years of musical training (χ^2^(1)=11.15, *p*=0.001). To help interpret these complex interactions, we conducted separate 2-way ANOVAs by group to assess the impact of music familiarity and condition on speech recognition in low vs. high PROMS listeners. High PROMS listeners’ comprehension was invariant to condition and familiarity effects (*ps*>0.065). However, the low PROMS group showed a familiarity*condition interaction (χ^2^(1)=3.63, *p*=0.042). Low musicality listeners showed poorer comprehension in music with vocals than without vocals but only during *unfamiliar* music (z=2.44, *p*=0.01). This vocal effect was not present for familiar music (z=2.20, *p*=0.69). When musicality was treated as continuous variable, there were no significant effects of any variable of interest on comprehension score (all *p*s > 0.311).

**Figure 4.**
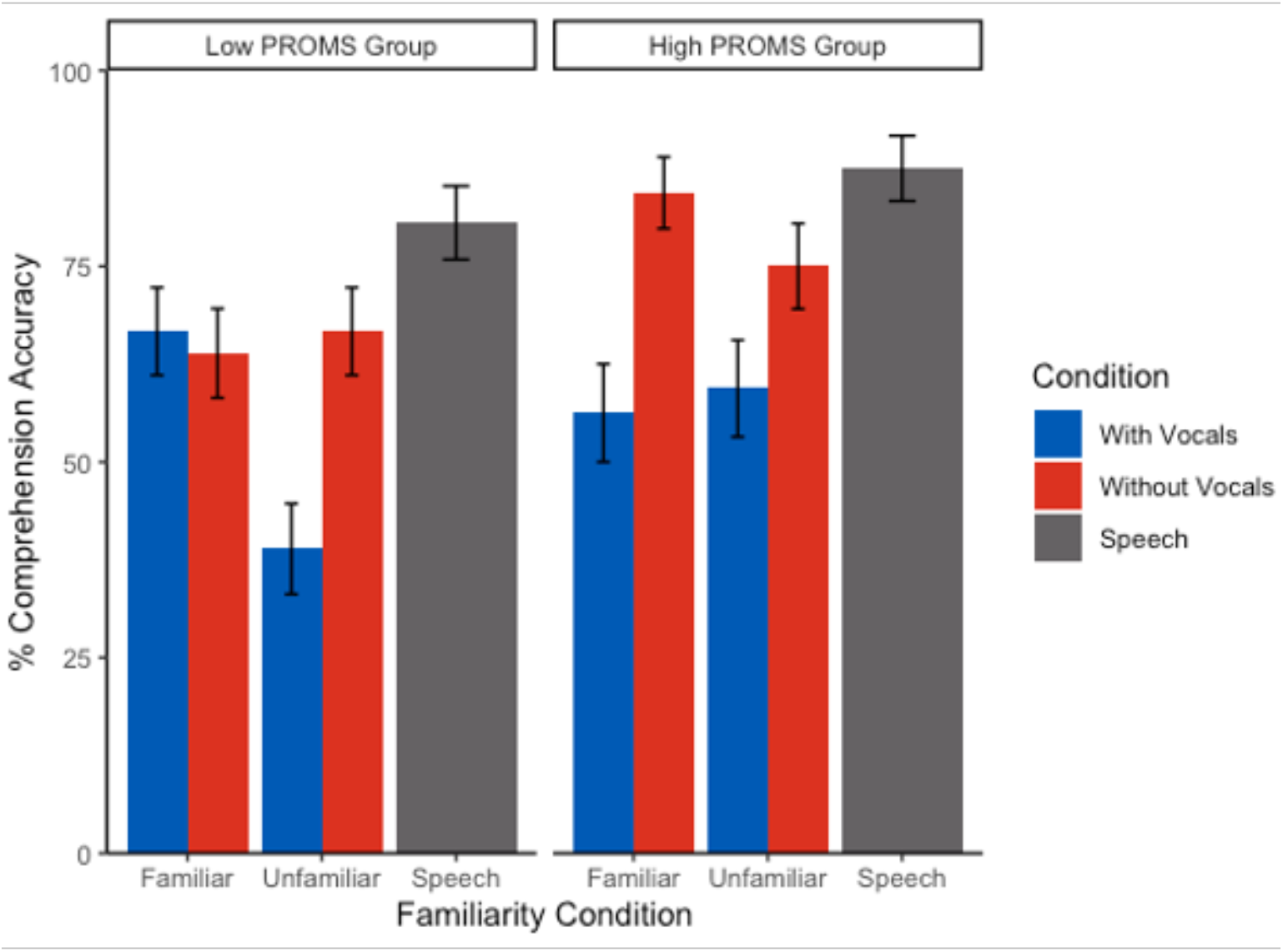
Speech comprehension performance varies with listeners’ musical skill level. Less musical listeners performed worse in music with vocals only in the unfamiliar music condition. However, more musical listeners did not show any effects of familiarity or song condition. PROMS = profile of music perception skills test. Error bars = ±1 s.e.m.

#### Neural TRFs

TRF latencies showed a significant interaction between familiarity and condition (*F*(1,45)=6.00, *p*=0.018, η^2^=0.12), where latencies for vocals were longer than instrumentals for only the unfamiliar music. There was no effect of musicality (*F*(1,15)=1.85, *p*=0.194). However, when treating musicality as a continuous variable, there was a familiarity*musicality interaction (*F*(1,45)=10.17, *p*=0.003, η^2^=0.18). TRF latencies were longer for *unfamiliar* music (*t*(45)=2.60, *p*=0.013) for the less musical listeners (i.e., lower PROMS scores) but were longer for *familiar* songs in higher PROMS scoring individuals (*t*(45)=2.63, *p*=0.012).

For TRF amplitudes, the omnibus ANOVA revealed an overall effect of condition (*F*(1,45)=4.88, *p*=0.032), where all listeners showed larger amplitudes for music with vocals (**Figure 5B**). There were no effects of familiarity or PROMS group (all *p*s > 0.1). There was also not an effect of musical training (*F*(1,15)=0.03, *p*=0.88), indicating again that the condition effect is not driven by experience. Treating musicality as a continuous variable showed no significant effects for any of the variables of interest (all *p*s >0.067).

**Figure 5.**
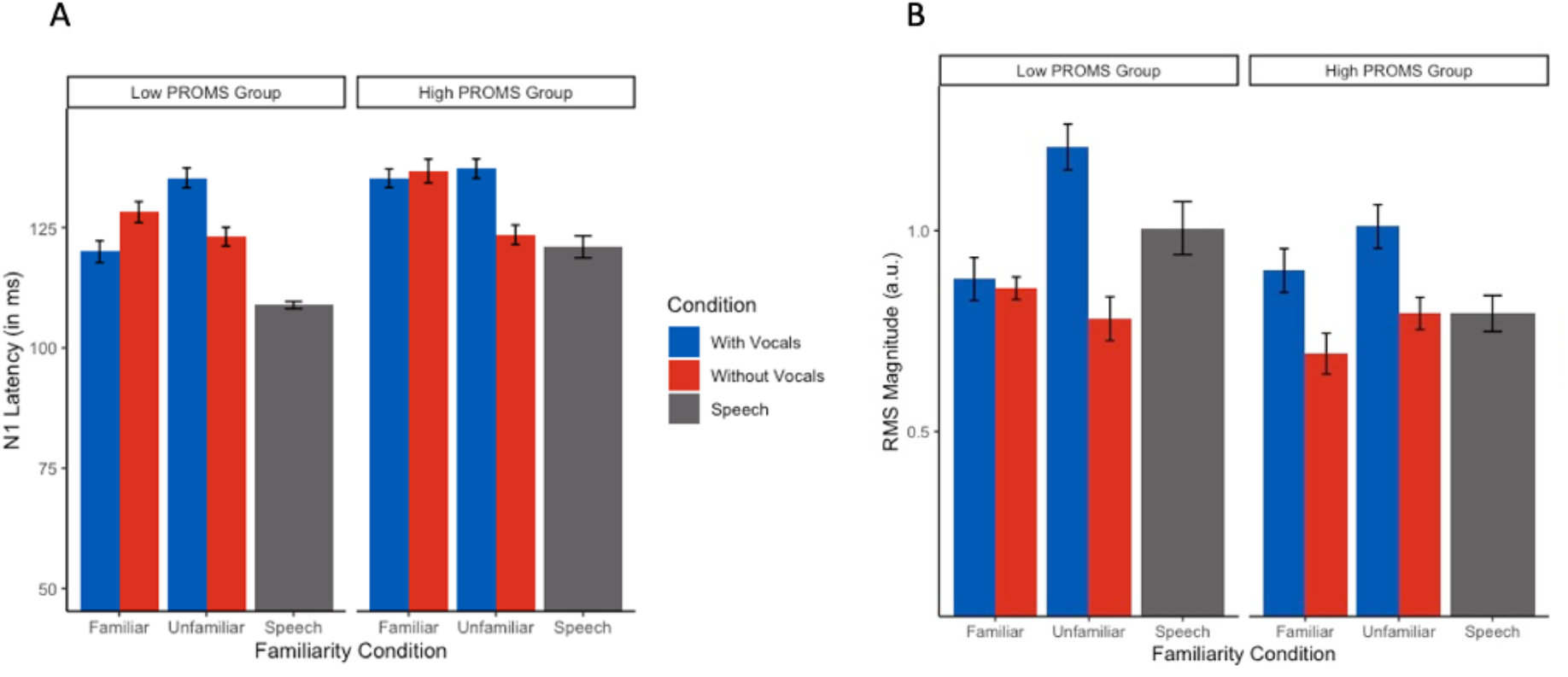
Speech TRFs vary as a function of musical aptitude (controlling for musical training). *(A)* Latencies were shorter for instrumental music than vocals only in the unfamiliar condition. (*B*) Instrumental music produced smaller speech responses than songs with vocals in both groups.

## DISCUSSION

As an innovative extension of the cocktail party problem, we compared speech comprehension and neural tracking of target speech amidst various *music* backdrops. We also manipulated the familiarity and vocals (i.e., song with lyrics or only instrumentals) of the music to evaluate how content and listeners’ familiarity of concurrent music backgrounds affect concurrent speech perception and its neural processing. Perception and neural encoding of speech was worse during music with vocals than purely instrumentals. However, more interestingly, the impact of vocals on speech coding varied based on the (un)familiarity of the background music. These findings indicate the monitoring of speech concurrent with music containing vocals might be more challenging for unfamiliar tunes. Our data suggest it is more difficult (i.e., harder for the brain to suppress the music masker) during certain types (unfamiliar, vocal) of music backdrops, likely through increased susceptibility to linguistic interference and/or misallocation of attention between speech and music streams. Moreover, these effects were exacerbated when accounting for musicality, suggesting listeners’ inherent auditory skills also impact their cocktail party speech processing.

In accordance with our previous behavioral study (Brown & Bidelman, 2022), we found speech comprehension was further impaired in music with vocals than in instrumental music. We attribute this decline to the informational masking introduced by the linguistic content of the vocals (Brouwer et al., 2021; Kidd & Colburn, 2017; Scharenborg & Larson, 2018). However, we previously found that comprehension was *worse* in unfamiliar music. If unfamiliar music is more distracting, suppression of the distractor (the background music) is impaired, thus decreasing the available cognitive resources needed to focus on the target speech (Lavie et al., 2004). Though the two studies used the same music maskers, the task here was more complex, assessing speech comprehension rather than target word recognition (as in Brown & Bidelman, 2022). These differences may contribute to the contradictory findings on familiarity. Indeed, the specific nature of the task can yield differential effects of music on concurrent speech processing, sometimes with opposite directions of effects (Brown & Bidelman, 2022; De Groot & Smedinga, 2014; Dong et al., 2022; Feng & Bidelman, 2015; Russo & Pichora-Fuller, 2008).

At the neural level, TRFs showed that brain tracking of speech was modulated by both familiarity and song condition. Overall, the presence of music prolonged TRF latencies to speech indicative of well-known masking effects observed in previous auditory EEG studies (Bidelman & Howell, 2016; Muncke et al., 2022). More critically, we found that condition and familiarity had an interactive effect on speech-evoked TRFs; neural latencies were strongly modulated by vocals relative to instrumental music but only for *unfamiliar* music. Responses were also larger during concurrent vocal compared to instrumental music. Intuitively, larger evoked responses are typically associated with a stronger representation or encoding of the speech signal (i.e., “bigger is better”) (Key et al., 2005). However, we found here that songs with vocals showed *worse* comprehension and longer neural latencies in conjunction with larger N1 responses. Indeed, TRF latencies were negatively correlated with speech recognition performance. It is well-established that larger N1 amplitudes are a marker of increased attention (Hillyard et al., 1973), analogous to the M100 peak in TRFs derived from neuromagnetic recordings (Ding & Simon, 2012a). Thus, the larger N1 responses to target speech we find in these difficult music conditions may reflect attentional load due to the increased listening demand of parsing speech from concurrent (especially unfamiliar) music. Larger N1 may also reflect increased listening effort (Strauss et al., 2010). Indeed, overexaggerated N1 to speech is also indicative of increased listening effort during speech processing, as observed in older adults with cognitive impairments (Bidelman et al., 2017). It is possible that neural speech tracking strength changed with attention and/or listening effort, where more distracting music maskers (i.e., unfamiliar/vocal songs) led to worse suppression of the masker, thus making it more difficult to maintain focus on the target. The stronger effects on speech processing we observe during unfamiliar vocal music might therefore reflect the influences of selective attention (Hillyard et al., 1973), with increased effort in maintaining that attention in more difficult listening conditions. This may also be comparable to a recent study that found a larger N400 during reading comprehension masked by music, reflecting increased semantic processing effort (Du et al., 2020). Still, the latency of neural effects observed here (~100 ms) suggest music challenges speech perception much earlier in the processing hierarchy.

Interestingly, we show that speech tracking at the “cocktail party” varies depending on the inherent musical skills of the listener. We found that low PROMS listeners were more impacted by unfamiliar music than the high PROMS group. Several studies have showed that more familiar music facilitates concurrent linguistic tasks by increasing arousal (Weiss et al., 2016) and generating expectancies (Bendixen, 2014; Russo & Pichora-Fuller, 2008). Musical ability is associated not only with speech-in-noise processing advantages (Hennessy et al., 2022; Mankel & Bidelman, 2018; Yoo & Bidelman, 2019), but also parsing complex auditory scenes (Johnson et al., 2021; Zendel & Alain, 2009). Indeed, there is some evidence that trained musicians also more successfully deploy attention in auditory and even non-auditory perceptual tasks (Bialystok & DePape, 2009; Román-Caballero et al., 2020; Yoo & Bidelman, 2019), including those related to cocktail party listening (Bidelman & Yoo, 2020). Moreover, non-musicians are more susceptible to informational and linguistic masking (Bidelman & Yoo, 2020; Oxenham et al., 2003)—though see Madsen et al. (2019). Here, low PROMS listeners may have been more distracted by the unfamiliar background music as a more challenging listening condition, which then becomes exacerbated by the presence of vocals. In this sense, less musical listeners (i.e., those with poorer auditory perceptual skills) might experience increased informational masking compared to their more musical peers. Our findings thus support notions that musical ability impacts cocktail party speech listening and one’s susceptibility to informational masking (Bidelman & Yoo, 2020; Oxenham et al., 2003; Swaminathan et al., 2015) but extend prior work by demonstrating such effects are not necessarily attributable to musicianship, *per se*, but instead depend on inherent listening abilities.

It is of note that most prior studies used years of formal music training (self-reported) as a metric for defining musicians and non-musicians, while we solely used aptitude scores. Though high- and low-PROMS groups were separable based on their years of music training, musicality group differences remained significant while controlling for training, meaning that the effects found in this study likely results from some combination of experience and natural auditory skills. Our data are consistent with emerging notions that listeners’ inherent rather than acquired musicality, *per se*, affect their speech-in-noise processing abilities (Mankel et al., 2020; Mankel & Bidelman, 2018).

In summary, our combined behavioral and neuroimaging results demonstrate that speech tracking is negatively affected both by familiarity and the presence of vocals in concurrent music. Furthermore, we show that these effects are modulated by musical ability, whereby less-musical listeners are more susceptible to these different background music characteristics. By using naturalistic, continuous stimuli, we simulated a realistic listening scenario, thus further adding to our understanding of the cocktail party phenomenon. Our findings also qualify prior studies (e.g., Dabaghi Varnosfaderani et al., 2021; Thompson et al., 2011) by suggesting that in addition to general arousal, familiarity and internal structure of music (e.g., presence or absence of vocals) might affect concurrent cognitive-linguistic processing.

Data are available on request from the authors.

## Acknowledgements

Work supported by the National Institute on Deafness and Other Communication Disorders (R01DC016267). Requests for data and materials should be directed to G.M.B.

## Author contributions

J.A.B. and G.M.B. designed the experiment, J.A.B. collected and analyzed the data, and J.A.B. and G.M.B. wrote the paper.

## References

Angel, L. A., Polzella, D. J., & Elvers, G. C. (2010). Background music and cognitive performance. In Perceptual and Motor Skills (Vol. 110, pp. 1059–1064).

Bates, D., Mächler, M., Bolker, B., & Walker, S. (2015). Fitting Linear Mixed-Effects Models Using lme4. Journal of Statistical Software, 67(1), 1–48. https://doi.org/10.18637/jss.v067.i01

Bendixen, A. (2014). Predictability effects in auditory scene analysis: A review. In Frontiers in Neuroscience (Vol. 8, pp. 1–16).

Bialystok, E., & DePape, A. M. (2009). Musical expertise, bilingualism, and executive functioning. In Journal of Experimental Psychology: Human Perception and Performance (2009/04/01 ed., Vol. 35, pp. 565–574). Department of Psychology, York University, Toronto, ON, Canada.

Bidelman, G. M., & Dexter, L. (2015). Bilinguals at the “cocktail party”: Dissociable neural activity in auditory-linguistic brain regions reveals neurobiological basis for nonnative listeners’ speech-in-noise recognition deficits. In Brain and Language (Vol. 143, pp. 32–41).

Bidelman, G. M., & Howell, M. (2016). Functional changes in inter-and intra-hemispheric auditory cortical processing underlying degraded speech perception. NeuroImage, 124, 581–590.

Bidelman, G. M., & Krishnan, A. (2010). Effects of reverberation on brainstem representation of speech in musicians and non-musicians. In Brain Research (Vol. 1355, pp. 112–125).

Bidelman, G. M., Lowther, J. E., Tak, S. H., & Alain, C. (2017). Mild cognitive impairment is characterized by deficient hierarchical speech coding between auditory brainstem and cortex. In Journal of Neuroscience (Vol. 37, pp. 3610–3620).

Bidelman, G. M., & Yoo, J. (2020). Musicians Show Improved Speech Segregation in Competitive, Multi-Talker Cocktail Party Scenarios. In Frontiers in Psychology (Vol. 11, pp. 1–11).

Brouwer, S., Akkermans, N., Hendriks, L., van Uden, H., & Wilms, V. (2021). “Lass frooby noo!” the interference of song lyrics and meaning on speech intelligibility. In Journal of Experimental Psychology: Applied: American Psychological Association (APA).

Brown, J. A., & Bidelman, G. M. (2022). Song properties and familiarity affect speech recognition in musical noise. In Psychomusicology: Music, Mind, and Brain.

Coffey, E. B. J., Mogilever, N. B., & Zatorre, R. J. (2017). Speech-in-noise perception in musicians: A review. In Hearing Research (Vol. 352, pp. 49–69): Elsevier B.V.

Crawford, H. J., & Strapp, C. M. (1994). Effects of vocal and instrumental music on visuospatial and verbal performance as moderated by studying preference and personality. In Personality and Individual Differences (Vol. 16, pp. 237–245).

Crosse, M. J., Di Liberto, G. M., Bednar, A., & Lalor, E. C. (2016). The multivariate temporal response function (mTRF) toolbox: A MATLAB toolbox for relating neural signals to continuous stimuli. In Frontiers in Human Neuroscience (Vol. 10).

Crosse, M. J., Zuk, N. J., Liberto, G. M. D., Nidiffer, A. R., Molholm, S., & Lalor, E. C. (2021). Linear Modeling of Neurophysiological Responses to Naturalistic Stimuli: Methodological Considerations for Applied Research. In Frontiers in Neuroscience (Vol. 15).

Dabaghi Varnosfaderani, S., Shahnazari, M., & Dabaghi, A. (2021). The influence of “happy” and “sad” background music on complexity, accuracy, and fluency of second-language speaking. Psychology of Music. https://doi.org/10.1177/03057356211033345

Darrow, A. A., Johnson, C., Agnew, S., Fuller, E. R., & Uchisaka, M. (2006). Effect of preferred music as a distraction on music majors’ and nonmusic majors’ selective attention. In Bulletin of the Council for Research in Music Education (pp. 21–31).

De Groot, A. M. B., & Smedinga, H. E. (2014). Let the music play!: A short-term but no long-term detrimental effect of vocal background music with familiar language lyrics on foreign language vocabulary learning. In Studies in Second Language Acquisition (Vol. 36, pp. 681–707).

Ding, N., Chatterjee, M., & Simon, J. Z. (2014). Robust cortical entrainment to the speech envelope relies on the spectro-temporal fine structure. NeuroImage, 88, 41–46. https://doi.org/10.1016/j.neuroimage.2013.10.054

Ding, N., & Simon, J. Z. (2012a). Emergence of neural encoding of auditory objects while listening to competing speakers. In Proceedings of the National Academy of Sciences of the United States of America (Vol. 109, pp. 11854–11859).

Ding, N., & Simon, J. Z. (2012b). Neural coding of continuous speech in auditory cortex during monaural and dichotic listening. In Journal of Neurophysiology (Vol. 107, pp. 78–89).

Dong, Y., Zheng, H.-Y., Wu, S. X.-Y., Huang, F.-Y., Peng, S.-N., Sun, S. Y.-K., & Zeng, H. (2022). The effect of Chinese pop background music on Chinese poetry reading comprehension. Psychology of Music, 1–22.

Du, M., Jiang, J., Li, Z., Man, D., & Jiang, C. (2020). The effects of background music on neural responses during reading comprehension. Sci Rep, 10(1), 18651. https://doi.org/10.1038/s41598-020-75623-3

Ekström, S. R., & Borg, E. (2011). Hearing speech in music. In Noise and Health (Vol. 13, pp. 277–285).

Eskridge, E., Galvin III, J., Aronoff, J., Li, T., & Fu, Q. J. (2012). Speech Perception with Music MAskers by Cochlear Implant Users and Normal Hearing Listeners. In Journal of Speech Language and Hearing Research (Vol. 55, pp. 800–810).

Feng, S., & Bidelman, G. M. (2015). Music listening and song familiarity modulate mind wandering and behavioral success during lexical processing. In Annual Meeting of the Cognitive science Society (CogSci 2015).

Forte, A. E., Etard, O., & Reichenbach, T. (2017). The human auditory brainstem response to running speech reveals a subcortical mechanism for selective attention. In bioRxiv.

Hennessy, S., Mack, W. J., & Habibi, A. (2022). Speech-in-noise perception in musicians and non-musicians: A multi-level meta-analysis. In Hearing Research (Vol. 416, pp. 108442): Elsevier B.V.

Hillyard, S. A., Hink, R. F., Schwent, V. L., & Picton, T. W. (1973). Electrical signs of selective attention in the human brain. In Science (Vol. 182, pp. 177–180).

Johnson, N., Shiju, A. M., Parmar, A., & Prabhu, P. (2021). Evaluation of auditory stream segregation in musicians and nonmusicians. In International Archives of Otorhinolaryngology (Vol. 25, pp. 77–80).

Johnsrude, I. S., Mackey, A., Hakyemez, H., Alexander, E., Trang, H. P., & Carlyon, R. P. (2013). Swinging at a cocktail party: Voice familiarity aids speech perception in the presence of a competing voice. In Psychological Science (2013/08/30 ed., Vol. 24, pp. 1995–2004). 1Department of Psychology, Queen’s University.

Kämpfe, J., Sedlmeier, P., & Renkewitz, F. (2011). The impact of background music on adult listeners: A meta-analysis. In Psychology of Music (Vol. 39, pp. 424–448).

Key, A. P. F., Dove, G. O., & Maguire, M. J. (2005). Linking brainwaves to the brain: An ERP primer. In Developmental Neuropsychology (Vol. 27, pp. 183–215).

Kidd, G., & Colburn, H. S. (2017). Informational Masking in Speech Recognition. In J. C. Middlebrooks, J. Z. Simon, A. N. Popper, & R. R. Fay (Eds.), The Auditory System at the Cocktail Party (pp. 75–109). Springer International Publishing. https://doi.org/10.1007/978-3-319-51662-2_4

Kulasingham, J. P., & Simon, J. Z. (2022). Algorithms for Estimating Time-Locked Neural Response Components in Cortical Processing of Continuous Speech. In bioRxiv (pp. 2022.2001.2018.476815).

Lalor, E. C., Power, A. J., Reilly, R. B., & Foxe, J. J. (2009). Resolving precise temporal processing properties of the auditory system using continuous stimuli. J Neurophysiol, 102(1), 349–359. https://doi.org/10.1152/jn.90896.2008

Lavie, N., Hirst, A., De Fockert, J. W., & Viding, E. (2004). Load theory of selective attention and cognitive control. In Journal of Experimental Psychology: General (Vol. 133, pp. 339–354).

Law, L. N., & Zentner, M. (2012). Assessing musical abilities objectively: construction and validation of the profile of music perception skills. PLoS One, 7(12), e52508. https://doi.org/10.1371/journal.pone.0052508

Lee, E. K., Lee, S. E., & Kwon, Y. S. (2020). The effect of lyrical and non-lyrical background music on different types of language processing - An ERP study. Korean Journal of Cognitive Science, 31(4), 155–178. https://doi.org/10.19066/cogsci.2020.31.4.003

MacCallum, R. C., Zhang, S., Preacher, K. J., & Rucker, D. (2002). On the practice of dichotomization of quantitative variables. In Psychological Methods (Vol. 7, pp. 19–40).

Madsen, S. M. K., Marschall, M., Dau, T., & Oxenham, A. J. (2019). Speech perception is similar for musicians and non-musicians across a wide range of conditions. In Scientific Reports (Vol. 9, pp. 10404).

Mankel, K., Barber, J., & Bidelman, G. M. (2020). Auditory categorical processing for speech is modulated by inherent musical listening skills. NeuroReport, 31(2), 162–166.

Mankel, K., & Bidelman, G. M. (2018). Inherent auditory skills rather than formal music training shape the neural encoding of speech. In Proceedings of the National Academy of Sciences (Vol. 115, pp. 13129–13134).

Martin, R. C., Wogalter, M. S., & Forlano, J. G. (1988). Reading comprehension in the presence of unattended speech and music. In Journal of Memory and Language (Vol. 27, pp. 382–398).

Muncke, J., Kuruvila, I., & Hoppe, U. (2022). Prediction of Speech Intelligibility by Means of EEG Responses to Sentences in Noise [Original Research]. Frontiers in Neuroscience, 16. https://doi.org/10.3389/fnins.2022.876421

Näätänen, R., & Picton, T. (1987). The N1 wave of the human electric and magnetic response to sound: A review and an analysis of the component structure. In Psychophysiology (1987/07/01 ed., Vol. 24, pp. 375–425).

Oldfield, R. (1971). The Assessment and Analysis of Handedness: The Edinburgh Inventory. In Neuropsychologia (Vol. 9, pp. 97–113).

Oostenveld, R., & Praamstra, P. (2001). The five percent electrode system for high-resolution EEG and ERP measurements. Clinical Neurophysiology, 112, 713–719.

Oxenham, A. J., Fligor, B. J., Mason, C. R., & Kidd Jr., G. (2003). Informational masking and musical training. In Journal of the Acoustical Society of America (2003/09/30 ed., Vol. 114, pp. 1543–1549). Research Laboratory of Electronics, Massachusetts Institute of Technology, Cambridge, Massachusetts 02139, USA.

Parbery-Clark, A., Skoe, E., & Kraus, N. (2009). Musical experience limits the degradative effects of background noise on the neural processing of sound. In Journal of Neuroscience (2009/11/13 ed., Vol. 29, pp. 14100–14107). Auditory Neuroscience Laboratory, Northwestern University, Evanston, Illinois 60208, USA.

Parbery-Clark, A., Skoe, E., Lam, C., & Kraus, N. (2009). Musician enhancement for speech-in-noise. In Ear and Hearing (2009/09/08 ed., Vol. 30, pp. 653–661).

Patston, L., & Tippett, L. (2011). The Effect of Background Music on Cognitive Performance in Musicians and Nonmusicians. In Musicians, Background Music, and Cognition (pp. 173–184).

Perham, N., & Currie, H. (2014). Does listening to preferred music improve reading comprehension performance?. In Applied Cognitive Psychology (Vol. 28, pp. 279–284).

Picton, T. W., Woods, D. L., & Proulx, G. B. (1978). Human auditory sustained potentials. I. The nature of the response. Electroencephalography and Clinical Neurophysiology, 45, 186–197.

Pisoni, D. B. (1993). Long-term memory in speech perception: Some new findings on talker variability, speaking rate and perceptual learning. In Speech Communication (Vol. 13, pp. 109–125).

Román-Caballero, R., Martín-Arévalo, E., & Lupiáñez, J. (2020). Attentional networks functioning and vigilance in expert musicians and non-musicians. In Psychological Research: Springer Berlin Heidelberg.

Russo, F. A., & Pichora-Fuller, M. K. (2008). Tune in or tune out: Age-related differences in listening to speech in music. In Ear and Hearing (Vol. 29, pp. 746–760).

Scharenborg, O., & Larson, M. (2018). Investigating the Effect of Music and Lyrics on Spoken-Word Recognition. In.

Souza, P., Gehani, N., Wright, R., & McCloy, D. (2013). The advantage of knowing the talker.

Strait, D. L., & Kraus, N. (2011). Can you hear me now? Musical training shapes functional brain networks for selective auditory attention and hearing speech in noise. In Frontiers in Psychology (2011/07/01 ed., Vol. 2, pp. 113). Auditory Neuroscience Laboratory, Northwestern University Evanston, IL, USA.

Strauss, D. J., Corona-Strauss, F. I., Trenado, C., Bernarding, C., Reith, W., Latzel, M., & Froehlich, M. (2010). Electrophysiological correlates of listening effort: Neurodynamical modeling and measurement. In Cognitive Neurodynamics (Vol. 4, pp. 119–131).

Swaminathan, J., Mason, C. R., Streeter, T. M., Best, V., Kidd Jr., G., & Patel, A. D. (2015). Musical training, individual differences and the cocktail party problem. In Scientific Reports (2015/06/27 ed., Vol. 5, pp. 11628). Department of Speech, Language and Hearing Sciences, Boston University, Boston, MA. Department of Psychology, Tufts University, Medford, MA.

Thompson, W. F., Schellenberg, E. G., & Letnic, A. K. (2011). Fast and loud background music disrupts reading comprehension. In Psychology of Music (Vol. 40, pp. 700–708).

Vasilev, M. R., Kirkby, J. A., & Angele, B. (2018). Auditory Distraction During Reading: A Bayesian Meta-Analysis of a Continuing Controversy. Perspect Psychol Sci, 13(5), 567–597. https://doi.org/10.1177/1745691617747398

Weiss, M. W., Trehub, S. E., Schellenberg, E. G., & Habashi, P. (2016). Pupils Dilate for Vocal or Familiar Music. In Journal of experimental psychology. Human perception and performance.

Yonan, C. A., & Sommers, M. S. (2000). The Effects of Talker Familiarity on Spoken Word Identification in Younger and Older Listeners. Psychology and Agjn.

Yoo, J., & Bidelman, G. M. (2019). Linguistic, perceptual, and cognitive factors underlying musicians’ benefits in noise-degraded speech perception. Hear Res, 377, 189–195. https://doi.org/10.1016/j.heares.2019.03.021

Zendel, B. R., & Alain, C. (2009). Concurrent sound segregation is enhanced in musicians. In Journal of Cognitive Neuroscience (2008/10/01 ed., Vol. 21, pp. 1488–1498). University of Toronto, Ontario, Canada.

